# Copy number variants implicate cardiac function and development pathways in earthquake-induced stress cardiomyopathy

**DOI:** 10.1101/144675

**Authors:** Cameron J. Lacey, Kit Doudney, Paul G. Bridgman, Peter M. George, Roger T. Mulder, Julie J. Zarifeh, Bridget Kimber, Murray J. Cadzow, Michael A. Black, Tony R. Merriman, Klaus Lehnert, Vivienne M Bickley, John F. Pearson, Vicky A. Cameron, Martin A. Kennedy

## Abstract

The pathophysiology of stress cardiomyopathy (SCM), also known as takotsubo syndrome, is poorly understood. SCM usually occurs sporadically, often in association with a stressful event, but clusters of cases are reported after major natural disasters. There is some evidence that this is a familial condition. We have examined three possible models for an underlying genetic predisposition to SCM. Our primary study cohort consists of 28 women who suffered SCM as a result of two devastating earthquakes that struck the city of Christchurch, New Zealand, in 2010 and 2011. To seek possible underlying genetic factors we carried out exome analysis, Cardio-MetaboChip genotyping array analysis and array comparative genomic hybridization on these subjects. The most striking finding from these analyses was the observation of a markedly elevated rate of rare, heterogeneous copy number variants (CNV) of uncertain clinical significance (in 12/28 subjects). Several of these CNVs clearly impacted on genes of cardiac relevance including *RBFOX1, GPC5, KCNRG, CHODL*, and *GPBP1L1*. There is no physical overlap between the CNVs, and the genes they impact do not fall into a clear pathophysiological pathway. However, the recognition that SCM cases display a high rate of unusual CNV, and that SCM predisposition may therefore be associated with these CNVs, offers a novel perspective and a new approach by which to understand this enigmatic condition.

## Introduction

Stress cardiomyopathy (SCM), also known as “broken heart syndrome” or takotsubo syndrome,^1; 2^ is a condition that captures widespread public interest. The cardiomyopathy is distinctive and the precipitating emotional event is typically clearly defined, however the mechanism for the cardiomyopathy and links between the psychological event and the physical illness are not understood.

Sporadic cases of SCM are estimated to account for 1-5% of acute coronary syndrome presentations.^3–5^ Predominantly the condition occurs in post-menopausal women,^6; 7^ and because of this, 5-10% of female presentations with suspected acute coronary syndrome are attributed to SCM.^8–11^ Although SCM can be fatal, the symptoms are commonly transient and patients generally have a good prognosis and recover well over a period of days to weeks.^12^ In classic descriptions the cardiomyopathy has a typical pattern but a number of variations are now widely recognised and it is increasingly apparent that cases can be quite heterogeneous.^13^

SCM occurring in clusters around the time of major disasters such as earthquakes, floods and bushfires is also well recognised.^6; 14–16^ Due to the large impact these events have upon hospital resources and medical infrastructure, it is rare for such clusters of SCM to be studied in any depth. This was made clear in reports from the Great East Japan Earthquake.^17^ In the Canterbury (New Zealand) earthquake sequence of 2010 and 2011 the two main events precipitated large case clusters of SCM.^17–22^ Unusually for a major natural disaster, the tertiary hospital in Christchurch continued to function, allowing the collection of a relatively large homogenous cohort of cases which have been followed over several years.^18–23^ Most research around this disorder has focused on sporadic SCM associated with heterogenous triggers.^4; 24–26^ Although the presentation of earthquake-associated SCM (EqSCM) appears to be similar to that of sporadic cases, a key difference is the homogenous nature of the trigger.

Various mechanisms have been postulated for takotsubo cardiomyopathy, including that the syndrome arises from stunning of the heart muscle (myocardium) as a result of either ischemia from spasm of the coronary arteries, or from the direct effect of catecholamines (dopamine, adrenaline or noradrenaline) on cardiac myocytes.^4; 24; 27; 28^ Despite suggestive pathophysiological observations and theories, most authors conclude that the aetiology of SCM is poorly understood, and we do not yet have satisfactory explanations for the origins of this condition.^27; 29–32 26; 32^ Some retrospective case series have suggested that the incidence of SCM is increased in patients with anxiety conditions, but in our studies we did not find any correlation with psychiatric or anxiety disorders.^19; 23^

Amongst the models that may be proposed for SCM aetiology, it is worth considering the possible contribution of genetic factors. Many forms of cardiomyopathy have genetic origins.^33; 34^ Hypertrophic cardiomyopathy is the most common form of familial heart disease and a leading cause of sudden cardiac death. It is inherited in an autosomal dominant Mendelian manner with variable expressivity and age-related penetrance.^33^ These cardiomyopathies show considerable genetic heterogeneity, with cases now attributed to some 1400 mutations in 11 genes, all of which contribute to cardiac sarcomere function. Familial dilated cardiomyopathy is also frequently attributable to an underlying genetic predisposition and at least 50 genes have now been implicated, with most eliciting disease as dominant mutations.^34^

Evidence for genetic contributions to SCM are not as strong as for other cardiomyopathies. However, there are several examples of familial occurrence of SCM involving siblings^35–37^ or mother-daughter pairs,^38–42^ and a large Swedish study of SCM identified three families in which several close relatives developed the condition.^43^ The overall rarity of SCM would suggest that these familial clusters are significant, and it is quite possible that more overt familial relationships in this disorder are obscured by the simultaneous requirement for two key circumstances (in most cases): post-menopausal status and environmental exposure to a sudden major stressful event. Occasional cases of SCM occur in younger women or males, and a proportion of patients report no preceding stressor,^44^ suggesting that an intrinsic pathogenic mechanism is involved. The recurrence of SCM in some patients, including one Christchurch EqSCM case,^22^ also implies a biological vulnerability.

These observations have prompted consideration of genetic susceptibility to this condition.^39; 42; 45; 46^ Until recently, genetic studies were restricted to candidate gene analysis in case series of sporadic SCM patients,^47–51^ but these have yielded mainly negative findings. One candidate gene study reported a significant difference in the frequency of a *GRK5* polymorphism in cases,^52^ but this has not been replicated and past history of single gene association studies suggests it is unlikely to be meaningful.^53^ More recently, another candidate gene study has implicated estrogen receptor genes as potential risk factors for SCM.^54^ In an effort to capture genome-wide data, exome sequencing^55–58^ was recently applied to a sample of sporadic SCM cases.^42^ Although this analysis did not reveal any difference in allele frequency or burden between SCM cases and population controls (28 adults with normal echocardiograms), it was noted that two thirds of the cases carried a rare deleterious variant within at least one gene of a large set of adrenergic pathway genes, and 11 genes harboured a variant in two or more cases. However, the significance of these rare variants remains unclear.

In this study, we set out to explore the role of genetic factors in predisposition to EqSCM. We specifically tested three discrete hypotheses for potential genetic contributions to risk of SCM: (i) an essentially Mendelian hypothesis that rare genetic variants in one or a few key genes cause predisposition, which was tested by whole exome sequencing (WES); (ii) that SCM was a complex disorder with genetic contributions from multiple common variants, which was tested using the Cardio-MetaboChip genotyping array; and (iii) that rare copy number variants (CNV) impacting on relevant genes contribute to risk, which was tested by array comparative genomic hybridization (aCGH).

## Material and Methods

### Cases

The September 2010 earthquake of magnitude 7.1 on the Richter scale (Mw 7.1) in Christchurch (New Zealand) triggered eight cases of EqSCM, and the shallow highly destructive quake (Mw 6.3) that followed in February 2011 triggered 21 cases over four days. One woman presented after both quakes ^22^, and one was in hospital during the initial quake. Enrolment of this latter participant was delayed, and her sample was available for aCGH analysis (n=28) but not for Cardio-Metabolome analysis (n=27). The steps leading to recruitment of our EqSCM cohort are detailed elsewhere^20; 21^, but briefly, our study commenced the day of the first earthquake with the creation of a register of prospectively identified earthquake stress cardiomyopathy cases. As our hospital was still functioning we could build a cohort with first-world data from complete single centre capture. After the second earthquake the study was extended.^20–22^

Inclusion criteria: i) Meeting modified Mayo criteria for stress cardiomyopathy and admitted to Christchurch Hospital within one week of either the September 2010 or February 2011 earthquake; ii) age over 18; iii) informed consent given. Exclusion criteria: i) unable to understand English sufficiently to be able to complete questionnaires.

All participants were recruited with informed consent, including discussion of the possibility of incidental findings from genetic analyses, and return of such findings after consultation with a medical geneticist. The Southern Health and Disability Ethics Committee (New Zealand) approved this study.

### DNA extraction

Peripheral blood samples were obtained from consenting participants. Genomic DNA was extracted from 3 mL peripheral blood using NucleoMag extraction kits (Machery-Nagel GmbH, Düren, Germany) on a KingFisher^™^ Flex Magnetic liquid-handling robot (Thermo Fisher Scientific, Inc, Waltham, MA). DNA was quantified by analysis with the Nanodrop™ (ThermoFisher), and, where appropriate, the Tapestation 4200 system (Agilent Technologies).

### Exome analysis

We applied WES to a subset (24 of 28) EqSCM cases. The exome capture and sequencing was carried out in two batches of 12, during 2012-13 (New Zealand Genomics Limited, Dunedin, New Zealand). DNA was processed with Illumina TruSeq sample preparation and exome enrichment kits (which capture ~62Mb of genomic DNA), and sequencing (100bp paired-end reads) was carried out on an Illumina HiSeq2000 system. Good quality sequence was obtained across all exomes, with very few unassigned reads, and greater than 20 million sequence reads per sample at mean quality scores (Phred) of Q37. Raw read data were aligned to the GRCh37 human reference genome using the Burrows-Wheeler Aligner (BWA),^59^ and processed through the Broad GATK pipeline.^60^ The alignment process included removal of reads from duplicate fragments, realignment around known indels, and recalibration of all base quality scores. Joint variant calling was performed with GATK’s HaplotypeCaller. This included *de novo* assembly at each potential variant locus. Variants were annotated and analysed using Ingenuity Variant Analysis (IVA) software (QIAGEN, Redwood City, CA, USA), MutationTaster2,^61^ SnpEff,^62^ SeattleSeq annotation server,^55^ and Galaxy (via usegalaxy.org).^63^ Allele frequencies and additional annotations were drawn from 1000 Genomes project,^64^ NHLBI GO Exome Sequencing Project (ESP), Seattle, WA (URL: http://evs.gs.washington.edu/EVS/), ClinVar,^65^ and Exome Aggregation Consortium (ExAC), Cambridge, MA (URL: http://exac.broadinstitute.org). Promising gene variants were inspected by Sanger sequence analysis on the appropriate genomic DNA samples.

### Cardio-MetaboChip Analysis

Three groups were genotyped using the Illumina Cardio-MetaboChip: 27 out of 28 female Christchurch EqSCM cases, 133 heart-healthy controls from the Canterbury Healthy Volunteers Study (HVOLs, 54 F / 79 M),^66^ and 157 patients recruited for an ongoing study of premature coronary heart disease and consented for genotyping (CHD, 64 F / 93 M). DNA samples were run on the Cardio-MetaboChip and scanned on the Illumina® iScan platform by AgResearch Limited (Invermay, New Zealand).

Quality control with summary analysis of allele and genotype frequencies, Hardy-Weinberg equilibrium tests, and missing genotype rates were performed with PLINK version 1.07 software.^67^ SNPs with a minor allele frequency of <0.05 and those that failed the Hardy-Weinberg equilibrium test (p<0.001) were excluded from the analysis, leaving 141,095 SNPs in the analysis (Table 1). Three samples from the CHD Study were also removed after analysis of relatedness. Principal Component Analysis (PCA, Eigenstrat 4.2) was performed on an independent subset of almost 50,000 SNPs; the first principal component explained 6% of the variation, subsequent components all less than 0.5%, and matched self-reported ethnicity (visual inspection). Hence the first principal component was subsequently included as a factor in the logistic regression. Logistic regression was performed to evaluate differences in SNP minor allele frequencies between groups, adjusted for ethnicity and gender, using an additive genetic model (R 3.01 software^68^). P values were adjusted for false discovery rate (FDR) using the Benjamini Yekuteli method.^69^ Pathway analysis was performed for the leading 100 SNPs in each pairwise group comparison, using MetaCore from GeneGo (Thomson Reuters).

**Table 1.** SNP markers removed from Cardio-MetaboChip analysis in quality control.

| Stage of analysis | Number of SNPs | Comments |
| --- | --- | --- |
| Assayed | 196,725 |  |
| No genotype | 10,927 |  |
| Mono allelic | 28,367 |  |
| MAF <0.004 | 44,424 | SNPs with only 1 minor allele found |
| Missing > 0.02 of samples | 36,884 |  |
| HWE fail | 130 | HWE_P <1e-20 and 1.5x more heterozygotes than expected |
| Total Removed | 65,622 | Note that SNPs could fail on two or more of the criteria above |
| **SNPs remaining in analysis** | **131,103** |  |

### CNV detection and analysis

Array comparative genomic hybridisation (aCGH) was undertaken on 28 EqSCM cases to examine structural variants in the cohort. For this analysis, we used either the Nimblegen 135k oligo array (CGX12) (Roche NimbleGen Inc, Madison, WI, USA), capable of genome-wide screening for CNV to a resolution of 10kb in well-categorised pathogenic genomic regions, and 50kb elsewhere or the Agilent 180k HD oligo array (Sureprint G3 Human 4x180k) (Agilent Technologies, Santa Clara, CA, USA), which has a similar resolution.

Pooled reference DNA samples (catalogue numbers G147A and G152A) were purchased from Promega (Madison, WI, USA). EqSCM case and reference DNA samples (0.5-1 μg each) were labelled with Cy3 and Cy5 dyes respectively, purified, hybridized, and washed according to Nimblegen and Agilent protocols. Microarrays were scanned on a GenePix 4000B laser scanner (Axon Instruments, CA, USA) or a G2600D Agilent SureScan microarray scanner (Agilent Technologies). Data was processed using Nimblescan (Roche Nimblegen Inc) or Cytogenomics software (Agilent Technologies) with the default algorithms and analysis settings, but with a 5 probe minimum calling threshold. All arrays passed QC metrics for derivative log ratio spread (DLRS) values of <0.2. CNV data was visualised and interpreted using Genoglyphix software (Perkin Elmer) and NCBI genome browser software (genome build hg19 (GRCh37)). The EqSCM aCGH data were assessed against many CNV databases including Genoglyphix Chromosome Aberration Database (containing over 14,000 validated variants from 50,000 samples) (Perkin Elmer, Waltham, MA, USA),^70^ DECIPHER,^71^ and the Database of Genomic Variation (DGV, containing CNV data from over 35,000 unaffected individuals).^72^ CNVs were classified as thought to be benign (TBB), uncertain clinical significance (UCS), or clinically significant (CS) using an evidence-based approach^73–76^ which included database comparisons (frequency in cases/controls and relation to phenotype), gene content, gene function and dosage sensitivity. A broad summary of our CNV interpretation algorithm is depicted in Figure S1. Rare CNVs are defined as those that occur at a frequency of ≤ 1%. We classified our rare CNV frequency using larger DGV studies containing >1000 individuals.^77–81^

## Results

### Exome Analysis

To test the potential for an essentially Mendelian predisposition to EqSCM, WES was carried out on 24 of the 28 Christchurch EqSCM cases. Several approaches to analysis of the identified variants were used, all of them hypothesising the involvement of gene variants with a low population minor allele frequency (MAF), that were over-represented in the EqSCM cohort. We carried out various iterations of filtering using variant allele frequency data derived from large population databases (1000 Genomes; NHBLI Exome Sequencing Project), followed by careful manual inspection of remaining variants. For example, excluding all variants present in these databases with an allele frequency > 3%, and selecting for any present in at least 4/24 EqSCM exomes, identified variants in 131 genes, none of which proved to be convincing on closer analysis. We also carried out ranking of gene variants by predicted functional impact using various approaches.^61; 62; 82^ Once again, none of the variants identified in these analyses proved to be significantly enriched amongst our EqSCM exomes.

Mitochondrial DNA reads can be recovered from exome data ^83^. We carried out manual inspection of BAM files of mitochondrial DNA for our exome data compared with non-disease control exomes ^84^. No unusual variants were detected in mitochondrial sequences of the EqSCM samples.

Finally, the 11 genes listed in Figure 2 of Goodloe et al (2014),^42^ as well as a gene recently proposed to play a role in SCM, *BAG3*,^46^ were carefully examined for presence of any rare variants in the EqSCM dataset. None of the previously identified variants,^42; 46^ and no other convincing rare variants in these genes, were detected.

### Cardio-MetaboChip Analysis

To test the possibility of a more complex, polygenetic basis to SCM risk, involving multiple variants of small effect size, Cardio-MetaboChip analysis was carried out. The Cardio-MetaboChip data for 27 EqSCM cases and 133 heart-healthy controls were compared by logistic regression (adjusted for ethnicity and gender, additive genetic model), first performed for pairwise comparisons across groups. No SNPs reached statistical significance of <0.05 after adjusting for false discovery rate (FDR) when comparing either the EqSCM and HVOLs, or the EqSCM and CHD samples. To investigate whether the top 100 of these SNPs mapped to gene pathways that might assist in understanding potential disease mechanisms underlying SCM, pathway analysis of the leading 100 SNPs in the EqSCM versus HVOLs pairwise comparison was performed in MetaCore. Disease Biomarker Pathway analysis identified Myocardial Ischemia as the third most enriched pathway (FDR-adjusted p=1.3e^-2^), featuring 11 SNP loci on our list out of 886 pathway objects, including annexin V, ANRIL, COL4A1, dynein, HXK4, nectin-2, PPAR-gamma, prolidase, Tcf(Lef), UGT, and VEGFR-2.

### aCGH Analysis

To test for potential involvement of CNVs in SCM, we applied aCGH to all cases. Of the 28 EqSCM cases examined by aCGH, twelve (42%) showed evidence of large, rare heterozygous CNVs classified as being of unclear clinical significance (Table 2), meaning that insufficient evidence is available for unequivocal determination of clinical significance.^73^ Of these CNVs, seven were deletions and six were duplications. All of the CNVs were different, and there was no physical overlap between the various CNVs. Each of these rare CNVs encompasses one or more genes, or their immediate upstream regulatory regions, and many of the genes included within the CNVs have functions of cardiac relevance. A full list of all CNVs detected in the cohort is presented in Table S1.

**Table 2.**
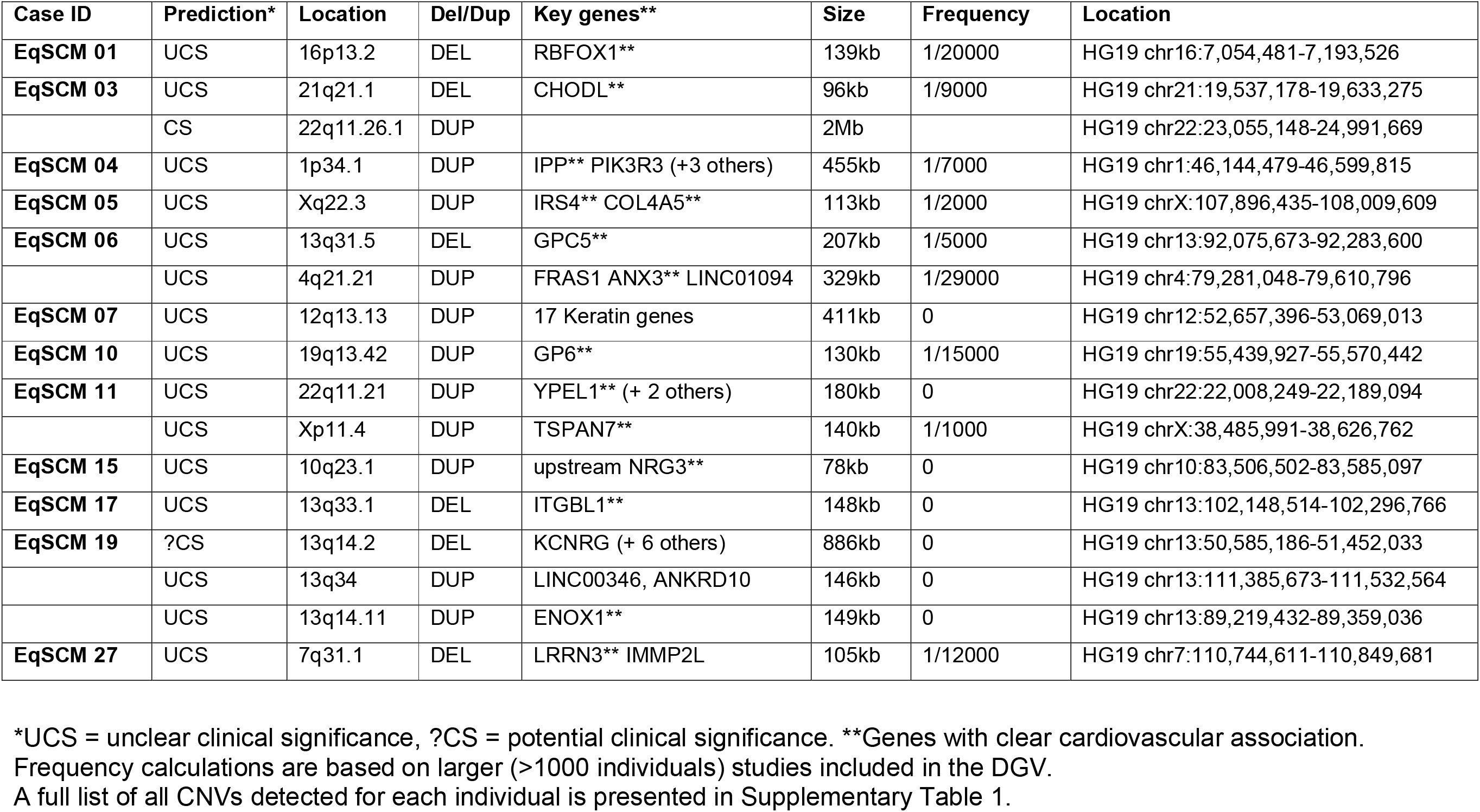
Rare CNVs detected in a cohort of 28 EqSCM cases.

| Case ID | Prediction* | Location | Del/Dup | Key genes** | Size | Frequency | Location |
| --- | --- | --- | --- | --- | --- | --- | --- |
| **EqSCM 01** | UCS | 16p13.2 | DEL | RBFOX1** | 139kb | 1/20000 | HG19 chr16:7,054,481-7,193,526 |
| **EqSCM 03** | UCS | 21q21.1 | DEL | CHODL** | 96kb | 1/9000 | HG19 chr21:19,537,178-19,633,275 |
|  | CS | 22q11.26.1 | DUP |  | 2Mb |  | HG19 chr22:23,055,148-24,991,669 |
| **EqSCM 04** | UCS | 1p34.1 | DUP | IPP** PIK3R3 (+3 others) | 455kb | 1/7000 | HG19 chr1:46,144,479-46,599,815 |
| **EqSCM 05** | UCS | Xq22.3 | DUP | IRS4** COL4A5** | 113kb | 1/2000 | HG19 chrX:107,896,435-108,009,609 |
| **EqSCM 06** | UCS | 13q31.5 | DEL | GPC5** | 207kb | 1/5000 | HG19 chr13:92,075,673-92,283,600 |
|  | UCS | 4q21.21 | DUP | FRAS1 ANX3** LINC01094 | 329kb | 1/29000 | HG19 chr4:79,281,048-79,610,796 |
| **EqSCM 07** | UCS | 12q13.13 | DUP | 17 Keratin genes | 411kb | 0 | HG19 chr12:52,657,396-53,069,013 |
| **EqSCM 10** | UCS | 19q13.42 | DUP | GP6** | 130kb | 1/15000 | HG19 chr19:55,439,927-55,570,442 |
| **EqSCM 11** | UCS | 22q11.21 | DUP | YPEL1** (+ 2 others) | 180kb | 0 | HG19 chr22:22,008,249-22,189,094 |
|  | UCS | Xp11.4 | DUP | TSPAN7** | 140kb | 1/1000 | HG19 chrX:38,485,991-38,626,762 |
| **EqSCM 15** | UCS | 10q23.1 | DUP | upstream NRG3** | 78kb | 0 | HG19 chr10:83,506,502-83,585,097 |
| **EqSCM 17** | UCS | 13q33.1 | DEL | ITGBL1** | 148kb | 0 | HG19 chr13:102,148,514-102,296,766 |
| **EqSCM 19** | ?CS | 13q14.2 | DEL | KCNRG (+ 6 others) | 886kb | 0 | HG19 chr13:50,585,186-51,452,033 |
|  | UCS | 13q34 | DUP | LINC00346, ANKRD10 | 146kb | 0 | HG19 chr13:111,385,673-111,532,564 |
|  | UCS | 13q14.11 | DUP | ENOX1** | 149kb | 0 | HG19 chr13:89,219,432-89,359,036 |
| **EqSCM 27** | UCS | 7q31.1 | DEL | LRRN3** IMMP2L | 105kb | 1/12000 | HG19 chr7:110,744,611-110,849,681 |

\*UCS = unclear clinical significance, ?CS = potential clinical significance. \*\*Genes with clear cardiovascular association. Frequency calculations are based on larger (>1000 individuals) studies included in the DGV. A full list of all CNVs detected for each individual is presented in Supplementary Table 1.

Three cases (EqSCM 01, 06 and 19) harboured deletions very likely to impact genes of high relevance to cardiomyopathy or cardiac function. In EqSCM 01, intragenic deletion of *RBFOX1* results in a single copy loss of one exon used by the majority of transcripts predicted for the gene (Figure 1). This exon contains the start methionine for the RBFOX1 protein, meaning the gene is most likely rendered non-functional. *RBFOX1* is an important RNA-binding protein mediating the incorporation of microexons into many transcripts associated with neurological patterning and tissue development,^85; 86^ particularly in the brain, heart and muscles. Intragenic deletions in *RBFOX1* have been observed in a range of conditions, including occasional cases with cardiac defects.^87; 88; 89^ Furthermore, RBFOX1-mediated RNA splicing was also recently shown to be an important regulator of cardiac hypertrophy and heart failure^90^.

**Figure 1.**
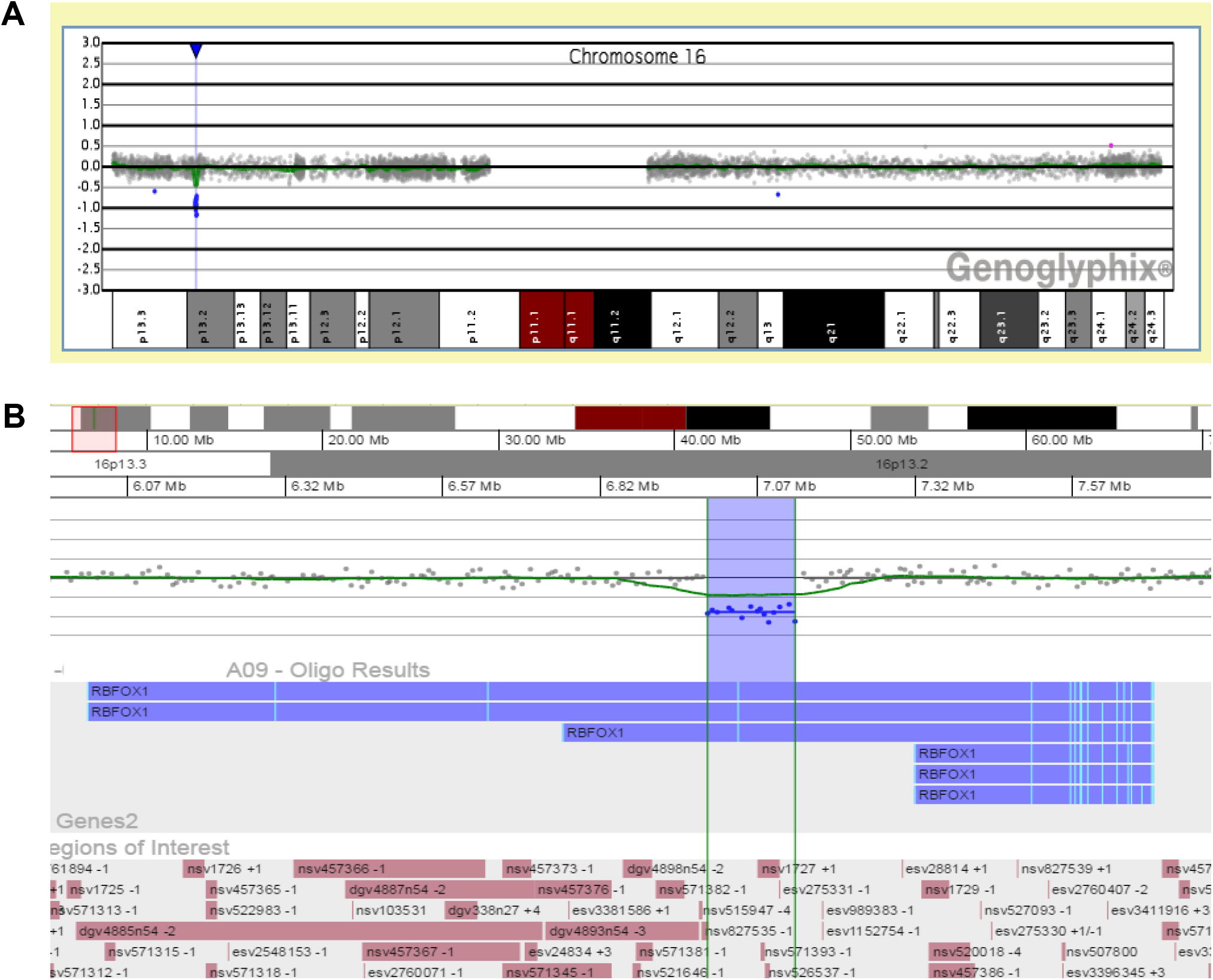
CNV detected in EqSCM case 01. **A:** Chromosomal location of the CNV at the RBFOX1 locus of chromosome 16. **B:** Enlargement of the fifteen probe deletion (139kb, delimited by vertical green lines and blue shading) illustrating loss of the fMet-containing exon (pale blue vertical bar) for three major RBFOX1 isoforms. DGV track of known CNVs shown at bottom of figure, beneath the genes2 and regions of interest tracks. Graphical views from Genoglyphix (PerkinElmer) software.

In the second case (EqSCM 06, Figure. 2), a heterozygous deletion encompassed exon 2 and the majority of intron 2 of the *Glypican 5 (GPC5)* locus. *GPC5* encodes a cell surface proteoglycan, which binds to the outer surface of the plasma membrane in the cardiovascular system and displays diverse functions including blood vessel formation after ischemic injury and proliferation of smooth muscle cells during atherogenesis.^91^ *GPC5* was also implicated by GWAS as a protective locus for sudden cardiac arrest,^92^ and other glypicans (GPC3, 4 and 6) have been associated with cardiac dysfunction.^93^ This case (EqSCM 06) also harbours a duplication on 4q21.21 involving the ANXA3 gene, which encodes a member of the annexin family, annexin A3. Members of this calcium-dependent phospholipid-binding protein family have a range of functions in the regulation of cellular growth and signal transduction pathways. Annexin A6 for example is the most abundant annexin expressed in the heart and its overexpression in mice has been shown to cause physiological alterations in contractility leading to dilated cardiomyopathy, while Annexin A6 knockout has been found to induce faster changes in Ca2+ transience and increased contractility.^94; 95^ Alterations in expression and activity of annexins A5 and A7 have also been found to be associated with regulation of Ca2+ handling in the heart.^96^ The function of annexin A3 is not fully understood, however it has been shown to play a role in endothelial migration and vascular development.^97^

The third case (EqSCM 19), contained a deletion at chr13q14.3. This region harbours at least 10 genes *(DLEU2, TRIM13, KCNRG, MIR16-1, MIR15A, DLEU1, DLEU1AS-1, ST13P4, DLEU7AS-1, DLEU7*, and *RNASEH2B-AS1)*, including several non-coding RNAs (DLEU genes and micro-RNA genes) and a gene *(KCNRG)* encoding a protein involved in the regulation of voltage-gated potassium channel activity. The micro-RNA genes mir-16-1 and mir-15a in this interval have been implicated in a range of cardiovascular phenotypes, including a role for mir-15a in postnatal mitotic arrest of cardiomyocytes.^98–100^ Two further chromosome 13 duplicated CNVs of approximately 150kb were classified as uncertain significance - one involving LINC00346 and ANKRD10 and the other containing ENOX1, postulated to affect vascular development based on zebrafish expression patterns^101^ (Table 2).

Beyond these three cases, cardiac or relevant neurological impacts appeared likely for many of the other rare CNVs identified in EqSCM cases (Table 2, Figure 2, Table S1), several of which are discussed below.

**Figure 2.**
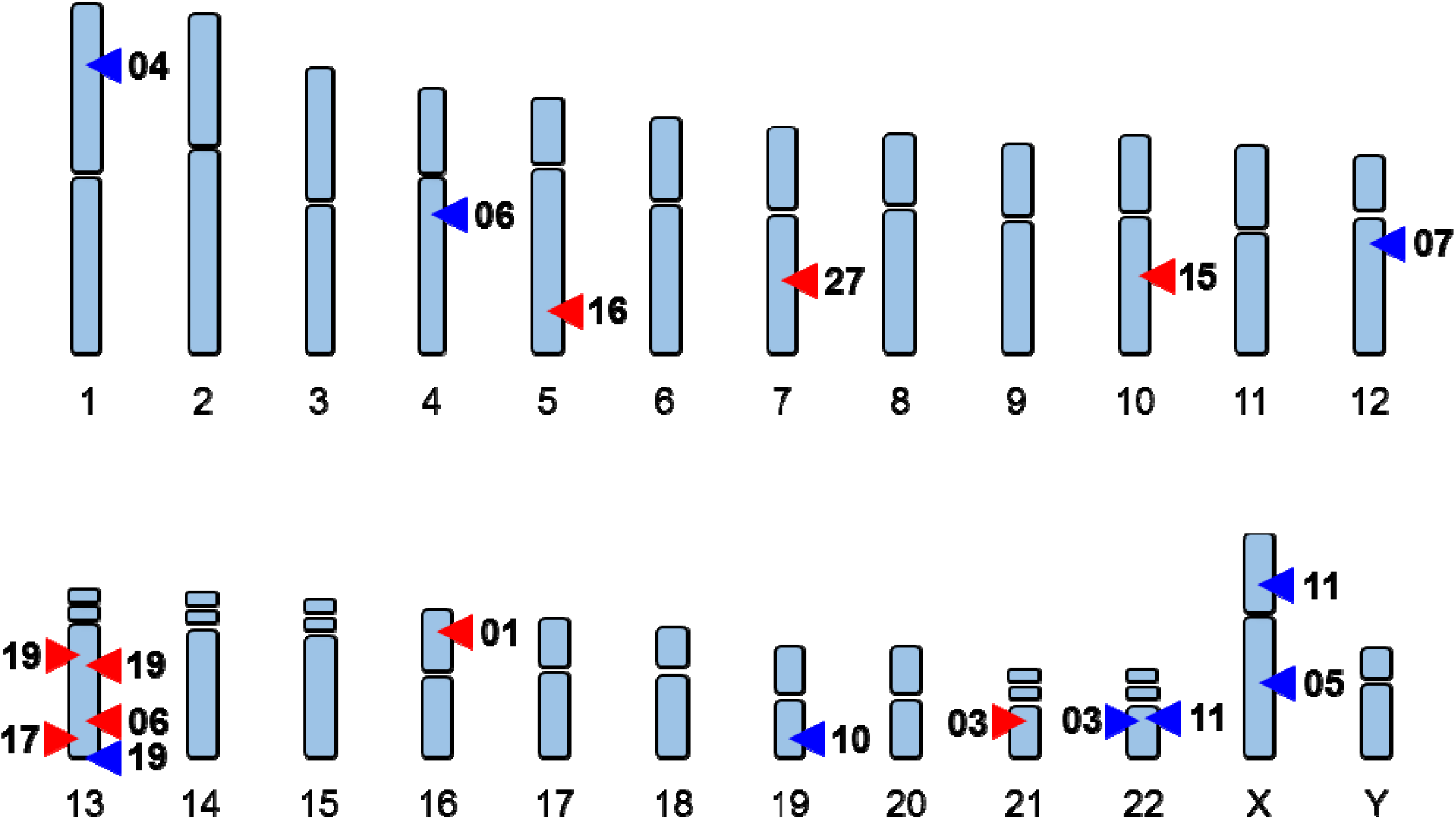
Genome wide distribution of CNVs. CNVs detected in 12 (of 28) EqSCM individuals by aCGH analysis. Numbers beside arrows relate to EqSCM patient number. Red arrows denote deletions, blue arrows duplications. Note that EqSCM 03, EqSCM 06, EqSCM 11 carry two rare CNVs, while EqSCM 19 contains three.

A 96kb heterozygous deletion in EqSCM 03 disrupts all predicted transcripts of the chondrolectin gene (*CHODL*), a membrane bound C-type lectin involved in muscle organ development, whose protein product is detected in heart and skeletal muscle by immunohistochemistry.^102^ In addition to this CNV, this patient carries a 1.94Mb duplication at 22q11.25, a locus containing 45 genes or miRNAs, associated with learning difficulties.^103; 104^ This individual, who exhibited a degree of cognitive impairment, had consented for clinically relevant findings to be forwarded to their General Practice clinician, who subsequently recommended genetic counselling for this individual.

A 455kb duplication within EqSCM 04 at 1p34.1 affects the genes *GPBP1L1, TMEM69, IPP, MAST2*, and *PIK3R3*. Smaller rare duplications in this region have been reported by the DGV database^105^ but none span the genes within this CNV. *GPBP1L1* is widely expressed in many tissues, including heart muscle^106^ and predicted to be involved in transcriptional regulation. *TMEM69*, a gene of unknown function, is most strongly expressed in heart tissue.^106^ IPP is a transcription factor with a 50 amino acid Kelch repeat known to interact with actin, while *MAST2* contains a PDZ domain and is another gene highly expressed in heart and skeletal muscle.^106^ The protein product of *PIK3R3*, phosphoinositide-3-kinase regulatory subunit 3, acts downstream of G-protein-coupled receptors in cardiac function,^107^ and is also a target for isoproterenol, which can trigger SCM-like conditions in humans and rodents.^12; 108–110^

Duplication of a long non-coding RNA (LOC101928358), the 3’ segment of *COL4A5* and the entire *IRS4* gene at Xq22.3 (9 probes, 113kb) was identified within case EqSCM 05. IRS4 is an insulin receptor molecule expressed in heart and skeletal muscle cells^111^ and other tissues such as brain, kidney and liver.^112^ Duplication of *IRS4* may be of functional significance, although copy number increases on the X chromosome of females may be counteracted to a degree by random X inactivation. Of note is a Genoglyphix Chromosome Aberration Database (GCAD^70^) case 52414 with a phenotype of low muscle tone, which has an identical duplication at this locus (as well as a 1p33 deletion). With regard to *IRS4*, Schreyer et al. (2003)^111^ found a more restricted tissue distribution than *IRS1* and *IRS2*, in primary human skeletal muscle cells and rat cardiac muscle and isolated cardiomyocytes. Although IRS4 protein function is still relatively unknown, the role of IRS proteins in general, acting as mediators of intracellular signalling from insulin and insulin-like growth factor 1 receptors, implicates IRS4 in cell growth and survival.^113^ It is interesting to note that PI 3-kinase (PI3K) signalling in HEH293T cells depends on IRS4, and that the IRS proteins relay signals from receptor tyrosine kinases to downstream components of signalling pathways,^111^ which we note is a connection with the *PIK3R3* gene duplicated in one of our other cases, EqSCM 04.

EqSCM 10 harboured a 130kb duplication of *NLRP7, NLRP2, GP6* and *RDH13* at 19q13.42. One similar DGV duplication has been seen in this area (nsv1062047^105^), but otherwise duplicated CNVs are generally much smaller and rare. *NLRP2* and *NLRP7* are genes that encode members of the NACHT, leucine rich repeat, and PYD containing (NLRP) protein family. These proteins are implicated in the activation of pro-inflammatory caspases. Recessive mutations ininin NLRP2/7 in humans are associated with reproductive disorders.^114^ Another gene in this duplicated cluster associated with disease is *GP6*, a platelet membrane glycoprotein, involved in collagen-induced platelet aggregation and thrombus formation, which is expressed at high levels in heart, kidney and whole blood.^106^ A *GP6* SNP (c.13254TC) has been implicated in recurrent cardiovascular events and mortality.^115^ Another study involving this SNP,^116^ found that hormone replacement therapy (HT) reduced the hazard ratio (HR) of CHD) events in patients with the *GP6* 13254TT genotype by 17% but increased the HR in patients with the TC+CC genotypes by 35% (adjusted interaction P < 0.001). The authors found that in postmenopausal women with established CHD, the *GP6* polymorphism, and another in *GP1B*, were predictors of CHD events and significantly modified the effects of HT on CHD risk.^116^

Duplication of *PPL2, YPEL1* and *MAPK1* on chromosome 22q11.21 (180kb) observed in EqSCM 11, does not appear in the DGV catalogue of CNVs in healthy individuals, and the consequences of overexpression of these genes, miRNAs or regulatory sequences are unknown. One of the affected genes, *MAPK1* (previously named *ERK* or *ERK2)*, may constitute a link to another kinase intracellular signalling pathway – the RAF-MEK-ERK kinase cascade, which in mice and human has an established role in the induction of cardiac tissue hypertrophy.^117^ Although not the kind of left ventricular enlargement seen in SCM, subtle copy number variation at *MAPK1* may influence signalling through this pathway. Another duplication in EqSCM 11 involving the 3’ half of *TSPAN7*, a member of the tetraspanin protein superfamily (Xp11.4) was noted as rare, and a similar (though larger) duplication was recently observed in a patient with Rolandic epilepsy.^118^

An agenic duplication 50kb upstream from *NRG3* (10q23.1) was seen in case EqSCM 15. This CNV could conceivably disrupt upstream regulatory regions of *NRG3*, which encodes an important ligand for the transmembrane tyrosine kinase receptor ERBB4. NRG3 has been shown to activate tyrosine phosphorylation of its cognate receptor, ERBB4, and is thought to influence neuroblast proliferation, migration and differentiation by signalling through ERBB4. *NRG3* is a strong candidate gene for schizophrenia, and neuregulin molecules and their receptors are involved in rat cardiac development and maintenance.^119^

Finally, the two rare deletions observed on chromosome 13 (13q21.33 and 13q33.1) in case EqSCM 17, fall into largely uncharacterised areas of the genome. The first is agenic, although there is a prediction of a spliced EST in the NCBI database, and the second occurs as two 50kb blocks within the *ITGBL1* gene. *ITGBL1* is most strongly expressed in aorta.^120; 121^

## Discussion

### Monogenic and polygenic models of risk

The Christchurch earthquakes repeatedly exposed the entire population of the city, approximately 350,000 people, to major stress and life disruption. Almost all patients presenting with EqSCM were post-menopausal females, consistent with other reports.^122^ We set out to explore three categories of genetic contributions to SCM predisposition, using WES to explore Mendelian models of risk, Cardio-MetaboChip analysis to test for polygenic risk factors, and aCGH analysis to evaluate the role of genomic structural variants.

Extensive analysis of the WES data did not yield any apparent enrichment of rare, damaging variants within exome regions amongst the EqSCM cases. Therefore, it seems unlikely that point mutations or small insertion-deletion (indels) in a single gene underlie predisposition to earthquake SCM. A limitation of this analysis is that it would have been unable to detect regulatory mutations, or other important variants, not included or well represented within the captured exome regions. Whole genome sequencing may therefore be warranted to further test the hypothesis of Mendelian underpinnings of SCM, as this approach could identify any regulatory variants not obtained with WES, and due to the absence of a DNA capture step, would also provide more uniform coverage of exons.

In a second approach, we explored the alternative hypothesis of polygenic risk alleles of small effect size using a case-control association study, with genotypes generated by the Cardio-MetaboChip. This chip allowed genotyping of ~200,000 SNPs previously identified through genome - wide association studies (GWAS) for risk of metabolic, atherosclerotic and cardiovascular diseases and traits.^123^ The traits covered by the panel of genetic variants on the chip include myocardial infarction (MI) and coronary heart disease (CHD), type 2 diabetes (T2D), T2D age diagnosed, T2D early onset, mean platelet volume, platelet count, white blood cell, HDL cholesterol, LDL cholesterol, triglycerides, total cholesterol, body mass index, waist hip ratio (BMI adjusted), waist circumference (BMI adjusted), height, percent fat mass, fasting glucose, fasting insulin, 2-hour glucose, HbA1c, systolic blood pressure, diastolic blood pressure and QT interval. This analysis did not yield variants of genome-wide significance in the SCM cases compared to either healthy controls or patients with coronary disease. Exploratory pathway analysis suggested that the EqSCM cases carried a greater burden of SNPs that mapped to a myocardial ischemia pathway compared to the healthy controls, although this must be interpreted with caution as our small sample set meant very limited statistical power. Two limitations of this analysis were the relatively constrained content of the Cardio-MetaboChip, which is less able to provide a rich dataset of genome-wide SNP genotypes than the chips commonly used for GWAS, and the relatively small cohort of cases available for study. Recruitment of a much larger SCM cohort with a view to a well-powered GWAS with a more extensive genotyping chip would therefore be a worthwhile future goal to more fully explore possible polygenic underpinnings of this disorder. We note the recent publication of a preliminary GWAS on 96 SCM cases and 475 healthy controls,^124^ and believe extension of this approach to larger cohorts is an important goal.

### Involvement of copy number variants

CNVs have been implicated in many diseases since the recognition a decade ago of their widespread distribution through the genome.^125–127^ Of note, rare CNVs are implicated in autism, epilepsy, schizophrenia, developmental delay and intellectual disability.^105; 128–132^ Cardiac conditions which involve CNVs include congenital left-sided heart disease,^133; 134^ congenital heart disease,^135; 136^ some cases of long QT syndrome,^137^ and Tetralogy of Fallot.^138^ Our final analysis, therefore, was to explore the potential involvement of CNV in risk of EqSCM, using aCGH analysis of all cases. Results from this analysis were striking, with 42% of EqSCM cases having a rare CNV of unclear clinical significance. The CNV detection rate for diagnostic aCGH in childhood developmental disorders such as autism, developmental delay and intellectual disabilities, is approximately 20-30%.^139–141^ A recent report of a large New Zealand aCGH case series (5,300 pre- and post-natal tests) reported CNVs in 28.3% of these clinically-selected cases.^142^ Our observation of a rate of 42% for the EqSCM case series is significantly greater (P < 0.02) than rates for the enriched case cohorts normally referred for clinical aCGH testing^139–142^. The CNVs detected in EqSCM cases were all different, and there were no physical overlaps between them. This situation is similar to the pattern of CNVs seen in other conditions, including rolandic epilepsies^118^ and congenital heart disease.^133; 135; 136^ Many of the CNVs we observed are likely to impact genes of potential relevance to physiological processes implicated in SCM.

We have taken a relatively conservative approach to categorising CNVs, in terms of rarity and predicted functional significance. For example, we did not include two CNVs located at 9p24.3, involving individuals EqSCM 14 and 28 - a deletion and duplication, respectively. These CNVs encompassed a region including the large isoform of *DOCK8* gene. *DOCK8* encodes a protein implicated in the regulation of the actin cytoskeleton,^143^ and *DOCK8* mutations cause autosomal recessive hyper-IgE syndrome.^144^ One reported case of a homozygous 129kb deletion in this region was associated with Graves’s disease and aortic aneurysm.^145^ However, several deletions of *DOCK8* are recorded in the DGV for unaffected individuals, therefore the CNVs in EqSCM 14 and 28 were categorised as TBB (thought to be benign). The approximately160kb duplication in EqSCM 28 was larger than the 44kb deletion in EqSCM 14, and it encompassed a second gene, *KANK1*. A small number of similar duplications have been recorded in the DGV, and therefore we did not consider this to be pathogenic. However, it is of interest that we see two relatively rare CNVs at this locus in our small EqSCM cohort.

Of the twelve EqSCM cases with rare CNVs, we consider that three (EqSCM 01, 06 and 19) contain CNVs that affect genes of high relevance to cardiomyopathy or cardiac function. The remaining candidate CNVs are also strong candidates with potential functional relevance. In one of our most highly-ranked candidate CNV containing cases (EqSCM 01) a large genomic deletion removes an exon of *RBFOX1* which contains the start codon used by the majority of transcripts predicted for the gene. A recent report by Gao *et al*. (2016) provided strong functional data that would support our hypothesis of this gene’s involvement as a susceptibility locus for SCM^90^. Their work with mouse models has shown RBFox1 deficiency in the heart promoted pressure overload–induced heart failure, and induction of *RBFox1* over-expression in these murine pressure-overload models, substantially attenuated cardiac hypertrophy and pathological manifestations^90^. The haploinsufficiency seen in EqSCM 01 at the *RBFOX1* locus may, in concert with other environmental or genetic factors, contribute to SCM through reduced global RNA splicing changes in the heart.

## Conclusion

Beginning with a cohort of 28 SCM cases triggered by two major earthquakes that caused extensive death and damage in Christchurch (New Zealand), we carried out exploratory analyses of three models for genetic predisposition to this disorder. Using WES and Cardio-MetaboChip genotyping analyses we did not detect an obvious role for exonic mutations in a monogenic model, or SNPs in a polygenic model, for SCM risk. However, our analysis of copy number variation in SCM cases revealed a high rate of occurrence of CNV categorised as of uncertain clinical significance. Most of the CNV we detected in SCM cases were rare, or not previously seen (Table 2).

These observations lead us to propose that SCM is a copy number variant disorder, whereby haploinsufficiency of genes overlapping deletions or over-expression of duplicated genes leads to relatively subtle modification of cardiac or adrenergic physiology, such that these individuals are at increased risk of suffering SCM when exposed to specific environmental triggers. Although no obvious single pathway relationships between the genes affected by these CNVs is apparent, most of the CNVs encompass loci relevant to cardiac function or cardioneuronal development.

In order to confirm whether SCM predisposition does indeed arise from CNVs, four key areas for future work need to be pursued. First, more widespread analysis is required of CNVs in many SCM cases. This would confirm whether our observation of a high rate of CNV in EqSCM also prevails in sporadic cases, and it will broaden the catalog of affected genes, helping to discern underlying signalling networks and physiological processes. In addition, with increasing numbers of cases, physical overlaps between CNVs in different individuals should become apparent, pinpointing key genomic regions for more intensive analysis. Second, the inheritance patterns of these CNVs must be established. It is unclear what proportion are *de novo* versus inherited from either parent. Third, there is a clear need for detailed physiological and gene expression analyses on appropriate cells, including cardiomyocytes, derived from SCM cases. Given the diversity of CNV seen in our SCM cases, this goal would most effectively be achieved by generation of induced-pluripotent stem cell (iPSC) lines from many patients and appropriate controls.^146; 147^ Finally, although our data implicate CNV as a significant genetic factor underlying SCM risk, it would seem wise to pursue an effective GWAS strategy to identify other genetic contributors to SCM and build on the initial study in this area.^124^ International initiatives to collate SCM cases^13^ should therefore ensure that consented DNA is available to provide appropriately large numbers of well phenotyped cases and controls to facilitate this goal.

Finally, we hope our observations implicating CNV in this unique case series of EqSCM will stimulate further studies of copy number variation in other SCM cohorts, and lead to an improved understanding of this perplexing and intriguing condition.

## Supplemental Data

Supplemental Data include one figure and one table and can be found with this article online at….

## Acknowledgements

This study was supported by an Earthquake research grant jointly awarded by the Health Research Council of New Zealand and Canterbury Medical Research Foundation. This study makes use of data generated by the DECIPHER community. A full list of centres who contributed to the generation of the data is available from http://decipher.sanger.ac.uk and via email from. Funding for the project was provided by the Wellcome Trust. We thank all participants for their agreement to be involved in this study, and Alan Aitchison and Allison Miller for preparation of DNA samples and Sanger sequencing analyses.

## Web Resources

1000 Genomes project, http://www.internationalgenome.org/

ClinVar, https://www.ncbi.nlm.nih.gov/clinvar/

Database of Genomic Variation (DGV), http://dgv.tcag.ca/dgv/app/home

DECIPHER, https://decipher.sanger.ac.uk/

Exome Aggregation Consortium (ExAC), http://exac.broadinstitute.org

Galaxy, https://galaxyproject.org/

GTex, https://www.gtexportal.org/home/

HUGO Gene Nomenclature Committee, http://www.genenames.org

MutationTaster2, http://www.mutationtaster.org/

NHLBI GO Exome Sequencing Project (ESP), https://esp.gs.washington.edu/drupal/

SeattleSeq annotation server, http://evs.gs.washington.edu/EVS/

## Supplemental data

Table S1: Complete CNV detection list of EqSCM cases in this study. (Separate Excel spreadsheet).

Figure S1: CNV interpretation algorithm (Separate word file doc)

## References

1. Dote, K., Sato, H., Tateishi, H., Uchida, T., and Ishihara, M. (1991). [Myocardial stunning due to simultaneous multivessel coronary spasms: a review of 5 cases]. J Cardiol 21, 203-214.

2. Tsuchihashi, K., Ueshima, K., Uchida, T., Oh-mura, N., Kimura, K., Owa, M., Yoshiyama, M., Miyazaki, S., Haze, K., Ogawa, H., et al. (2001). Transient left ventricular apical ballooning without coronary artery stenosis: a novel heart syndrome mimicking acute myocardial infarction. Angina Pectoris-Myocardial Infarction Investigations in Japan. Journal of the American College of Cardiology 38, 11-18.

3. Sharkey, S.W. (2013). Takotsubo cardiomyopathy: natural history. Heart Fail Clin 9, 123-136, vii.

4. Sharkey, S.W., Windenburg, D.C., Lesser, J.R., Maron, M.S., Hauser, R.G., Lesser, J.N., Haas, T.S., Hodges, J.S., and Maron, B.J. (2010). Natural history and expansive clinical profile of stress (tako-tsubo) cardiomyopathy. J Am Coll Cardiol 55, 333-341.

5. Eshtehardi, P., Koestner, S.C., Adorjan, P., Windecker, S., Meier, B., Hess, O.M., Wahl, A., and Cook, S. (2009). Transient apical ballooning syndrome--clinical characteristics, ballooning pattern, and long-term follow-up in a Swiss population. Int J Cardiol 135, 370-375.

6. Watanabe, H., Kodama, M., Okura, Y., Aizawa, Y., Tanabe, N., Chinushi, M., Nakamura, Y., Nagai, T., Sato, M., and Okabe, M. (2005). Impact of Earthquakes on Takotsubo Cardiomyopathy. Jama 294, 305-307.

7. Strunk, B., Shaw, R.E., Bull, S., Adams, J., Baer, M., Gershengorn, K., Kao, A., Keeffe, B., Sklar, J., Sperling, D., et al. (2006). High incidence of focal left ventricular wall motion abnormalities and normal coronary arteries in patients with myocardial infarctions presenting to a community hospital. The Journal of invasive cardiology 18, 376-381.

8. Sy, F., Basraon, J., Zheng, H., Singh, M., Richina, J., and Ambrose, J.A. (2013). Frequency of Takotsubo cardiomyopathy in postmenopausal women presenting with an acute coronary syndrome. Am J Cardiol 112, 479-482.

9. Eitel, I., von Knobelsdorff-Brenkenhoff, F., Bernhardt, P., Carbone, I., Muellerleile, K., Aldrovandi, A., Francone, M., Desch, S., Gutberlet, M., Strohm, O., et al. (2011). Clinical characteristics and cardiovascular magnetic resonance findings in stress (takotsubo) cardiomyopathy. JAMA 306, 277-286.

10. Prasad, A., Dangas, G., Srinivasan, M., Yu, J., Gersh, B.J., Mehran, R., and Stone, G.W. (2014). Incidence and angiographic characteristics of patients with apical ballooning syndrome (takotsubo/stress cardiomyopathy) in the HORIZONS-AMI trial: an analysis from a multicenter, international study of ST-elevation myocardial infarction. Catheter Cardiovasc Interv 83, 343-348.

11. Deshmukh, A., Kumar, G., Pant, S., Rihal, C., Murugiah, K., and Mehta, J.L. (2012). Prevalence of Takotsubo cardiomyopathy in the United States. American heart journal 164, 66-71 e61.

12. Wright, P.T., Tranter, M.H., Morley-Smith, A.C., and Lyon, A.R. (2014). Pathophysiology of takotsubo syndrome: temporal phases of cardiovascular responses to extreme stress. Circ J 78, 1550-1558.

13. Templin, C., Ghadri, J.R., Diekmann, J., Napp, L.C., Bataiosu, D.R., Jaguszewski, M., Cammann, V.L., Sarcon, A., Geyer, V., Neumann, C.A., et al. (2015). Clinical Features and Outcomes of Takotsubo (Stress) Cardiomyopathy. N Engl J Med 373, 929-938.

14. Steptoe, A. (2009). The impact of natural disasters on myocardial infarction. Heart 95, 1972-1973.

15. Butterly, S.J., Indrajith, M., Garrahy, P., Ng, A.C., Gould, P.A., and Wang, W.Y. (2013). Stress-induced takotsubo cardiomyopathy in survivors of the 2011 Queensland floods. Med J Aust 198, 109-110.

16. Leor, J., Poole, W.K., and Kloner, R.A. (1996). Sudden cardiac death triggered by an earthquake. N Engl J Med 334, 413-419.

17. Aoki, T., Takahashi, J., Fukumoto, Y., Yasuda, S., Ito, K., Miyata, S., Shinozaki, T., Inoue, K., Yagi, T., Komaru, T., et al. (2013). Effect of the Great East Japan Earthquake on cardiovascular diseases--report from the 10 hospitals in the disaster area. Circ J 77, 490-493.

18. Bridgman, P.G., Finsterer, J., Lacey, C., Kimber, B., Parkin, P.J., Miller, A.L., and Kennedy, M.A. (2016). CTG-repeat expansions in the DMPK gene do not cause takotsubo syndrome. Int J Cardiol 203, 107-108.

19. Lacey, C., Mulder, R., Bridgman, P., Kimber, B., Zarifeh, J., Kennedy, M., and Cameron, V. (2014). Broken heart syndrome - is it a psychosomatic disorder? J Psychosom Res 77, 158-160.

20. Chan, C., Elliott, J., Troughton, R., Frampton, C., Smyth, D., Crozier, I., and Bridgman, P. (2013). Acute myocardial infarction and stress cardiomyopathy following the Christchurch earthquakes. PLoS One 8, e68504.

21. Chan, C., Troughton, R., Elliott, J., Zarifeh, J., and Bridgman, P. (2014). One-year follow-up of the 2011 Christchurch Earthquake stress cardiomyopathy cases. N Z Med J 127, 15-22.

22. Bridgman, P.G., Chan, C.W., and Elliott, J.M. (2012). A case of recurrent earthquake stress cardiomyopathy with a differing wall motion abnormality. Echocardiography 29, E26-27.

23. Zarifeh, J.A., Mulder, R.T., Kerr, A.J., Chan, C.W., and Bridgman, P.G. (2012). Psychology of earthquake-induced stress cardiomyopathy, myocardial infarction and non-cardiac chest pain. Intern Med J 42, 369-373.

24. Wittstein, I.S. (2012). Stress cardiomyopathy: a syndrome of catecholamine-mediated myocardial stunning? Cell Mol Neurobiol 32, 847-857.

25. Coupez, E., Eschalier, R., Pereira, B., Pierrard, R., Souteyrand, G., Clerfond, G., Citron, B., Lusson, J.R., Mansencal, N., and Motreff, P. (2014). A single pathophysiological pathway in Takotsubo cardiomyopathy: Catecholaminergic stress. Arch Cardiovasc Dis 107, 245-252.

26. Wittstein, I.S. (2012). Stress cardiomyopathy: a syndrome of catecholamine-mediated myocardial stunning? Cell Mol Neurobiol 32, 847-857.

27. Wittstein, I.S., Thiemann, D.R., Lima, J.A., Baughman, K.L., Schulman, S.P., Gerstenblith, G., Wu, K.C., Rade, J.J., Bivalacqua, T.J., and Champion, H.C. (2005). Neurohumoral features of myocardial stunning due to sudden emotional stress. N Engl J Med 352, 539-548.

28. Nef, H.M., Mollmann, H., Kostin, S., Troidl, C., Voss, S., Weber, M., Dill, T., Rolf, A., Brandt, R., Hamm, C.W., et al. (2007). Tako-Tsubo cardiomyopathy: intraindividual structural analysis in the acute phase and after functional recovery. Eur Heart J 28, 2456-2464.

29. Prasad, A., Lerman, A., and Rihal, C.S. (2008). Apical ballooning syndrome (Tako-Tsubo or stress cardiomyopathy): a mimic of acute myocardial infarction. Am Heart J 155, 408-417.

30. Summers, M.R., Lennon, R.J., and Prasad, A. (2010). Pre-morbid psychiatric and cardiovascular diseases in apical ballooning syndrome (tako-tsubo/stress-induced cardiomyopathy): potential pre-disposing factors? J Am Coll Cardiol 55, 700-701.

31. Wright, P.T., Tranter, M.H., Morley-Smith, A.C., and Lyon, A.R. (2014). Pathophysiology of takotsubo syndrome. Circ J 78, 1550-1558.

32. Nef, H.M., Mollmann, H., Akashi, Y.J., and Hamm, C.W. (2010). Mechanisms of stress (Takotsubo) cardiomyopathy. Nat Rev Cardiol 7, 187-193.

33. Maron, B.J., Maron, M.S., and Semsarian, C. (2012). Genetics of hypertrophic cardiomyopathy after 20 years: clinical perspectives. J Am Coll Cardiol 60, 705-715.

34. McNally, E.M., Golbus, J.R., and Puckelwartz, M.J. (2013). Genetic mutations and mechanisms in dilated cardiomyopathy. J Clin Invest 123, 19-26.

35. Musumeci, B., Saponaro, A., Pagannone, E., Proietti, G., Mastromarino, V., Conti, E., Tubaro, M., Volpe, M., and Autore, C. (2013). Simultaneous Takotsubo syndrome in two sisters. Int J Cardiol 165, e49-50.

36. Ikutomi, M., Yamasaki, M., Matsusita, M., Watari, Y., Arashi, H., Endo, G., Yamaguchi, J., and Ohnishi, S. (2014). Takotsubo cardiomyopathy in siblings. Heart Vessels 29, 119-122.

37. Pison, L., De Vusser, P., and Mullens, W. (2004). Apical ballooning in relatives. Heart 90, e67.

38. Sharkey, S.W., Lips, D.L., Pink, V.R., and Maron, B.J. (2013). Daughter-mother tako-tsubo cardiomyopathy. Am J Cardiol 112, 137-138.

39. Cherian, J., Angelis, D., Filiberti, A., and Saperia, G. (2007). Can takotsubo cardiomyopathy be familial? Int J Cardiol 121, 74-75.

40. Kumar, G., Holmes, D.R., Jr., and Prasad, A. (2010). “Familial” apical ballooning syndrome (Takotsubo cardiomyopathy). Int J Cardiol 144, 444-445.

41. Subban, V., Ramachandran, S., Victor, S.M., Gnanaraj, A., and Ajit, M.S. (2012). Apical ballooning syndrome in first degree relatives. Indian Heart J 64, 607-609.

42. Goodloe, A.H., Evans, J.M., Middha, S., Prasad, A., and Olson, T.M. (2014). Characterizing genetic variation of adrenergic signalling pathways in Takotsubo (stress) cardiomyopathy exomes. Eur J Heart Fail 16, 942-949.

43. Schultz, T., Shao, Y., Redfors, B., Sverrisdottir, Y.B., Ramunddal, T., Albertsson, P., Matejka, G., and Omerovic, E. (2012). Stress-induced cardiomyopathy in Sweden: evidence for different ethnic predisposition and altered cardio-circulatory status. Cardiology 122, 180-186.

44. Pilgrim, T.M., and Wyss, T.R. (2008). Takotsubo cardiomyopathy or transient left ventricular apical ballooning syndrome: A systematic review. Int J Cardiol 124, 283-292.

45. Limongelli, G., D’Alessandro, R., Masarone, D., Maddaloni, V., Vriz, O., Minisini, R., Citro, R., Calabro, P., Russo, M.G., Calabro, R., et al. (2013). Takotsubo cardiomyopathy: do the genetics matter? Heart Fail Clin 9, 207-216, ix.

46. d’Avenia, M., Citro, R., De Marco, M., Veronese, A., Rosati, A., Visone, R., Leptidis, S., Philippen, L., Vitale, G., Cavallo, A., et al. (2015). A novel miR-371a-5p-mediated pathway, leading to BAG3 upregulation in cardiomyocytes in response to epinephrine, is lost in Takotsubo cardiomyopathy. Cell death & disease 6, e1948.

47. Figtree, G.A., Bagnall, R.D., Abdulla, I., Buchholz, S., Galougahi, K.K., Yan, W., Tan, T., Neil, C., Horowitz, J.D., Semsarian, C., et al. (2013). No association of G-protein-coupled receptor kinase 5 or beta-adrenergic receptor polymorphisms with Takotsubo cardiomyopathy in a large Australian cohort. Eur J Heart Fail 15, 730-733.

48. Sharkey, S.W., Maron, B.J., Nelson, P., Parpart, M., Maron, M.S., and Bristow, M.R. (2009). Adrenergic receptor polymorphisms in patients with stress (tako-tsubo) cardiomyopathy. J Cardiol 53, 53-57.

49. Handy, A.D., Prasad, A., and Olson, T.M. (2009). Investigating genetic variation of adrenergic receptors in familial stress cardiomyopathy (apical ballooning syndrome). Journal of cardiology 54, 516-517.

50. Spinelli, L., Trimarco, V., Di Marino, S., Marino, M., Iaccarino, G., and Trimarco, B. (2010). L41Q polymorphism of the G protein coupled receptor kinase 5 is associated with left ventricular apical ballooning syndrome. European journal of heart failure 12, 13-16.

51. Vriz, O., Minisini, R., Citro, R., Guerra, V., Zito, C., De Luca, G., Pavan, D., Pirisi, M., Limongelli, G., and Bossone, E. (2011). Analysis of beta1 and beta2-adrenergic receptors polymorphism in patients with apical ballooning cardiomyopathy. Acta Cardiol 66, 787-790.

52. Novo, G., Giambanco, S., Guglielmo, M., Arvigo, L., Sutera, M.R., Giambanco, F., Evola, S., Vaccarino, L., Bova, M., Lio, D., et al. (2014). G-protein-coupled receptor kinase 5 polymorphism and Takotsubo cardiomyopathy. J Cardiovasc Med (Hagerstown).

53. Munafo, M.R., and Flint, J. (2004). Meta-analysis of genetic association studies. Trends Genet 20, 439-444.

54. Pizzino, G., Bitto, A., Crea, P., Khandheria, B., Vriz, O., Carerj, S., Squadrito, F., Minisini, R., Citro, R., Cusma-Piccione, M., et al. (2017). Takotsubo syndrome and estrogen receptor genes: partners in crime? J Cardiovasc Med (Hagerstown) 18, 268-276.

55. Ng, S.B., Turner, E.H., Robertson, P.D., Flygare, S.D., Bigham, A.W., Lee, C., Shaffer, T., Wong, M., Bhattacharjee, A., Eichler, E.E., et al. (2009). Targeted capture and massively parallel sequencing of 12 human exomes. Nature 461, 272-276.

56. Bamshad, M.J., Ng, S.B., Bigham, A.W., Tabor, H.K., Emond, M.J., Nickerson, D.A., and Shendure, J. (2011). Exome sequencing as a tool for Mendelian disease gene discovery. Nature reviews Genetics 12, 745-755.

57. Yang, Y., Muzny, D.M., Reid, J.G., Bainbridge, M.N., Willis, A., Ward, P.A., Braxton, A., Beuten, J., Xia, F., Niu, Z., et al. (2013). Clinical whole-exome sequencing for the diagnosis of mendelian disorders. N Engl J Med 369, 1502-1511.

58. Chugh, S.S., and Huertas-Vazquez, A. (2014). Inherited arrhythmia syndromes: exome sequencing opens a new door to diagnosis. J Am Coll Cardiol 63, 267-268.

59. Li, H., and Durbin, R. (2009). Fast and accurate short read alignment with Burrows-Wheeler transform. Bioinformatics 25, 1754-1760.

60. McKenna, A., Hanna, M., Banks, E., Sivachenko, A., Cibulskis, K., Kernytsky, A., Garimella, K., Altshuler, D., Gabriel, S., Daly, M., et al. (2010). The Genome Analysis Toolkit: a MapReduce framework for analyzing next-generation DNA sequencing data. Genome research 20, 1297-1303.

61. Schwarz, J.M., Cooper, D.N., Schuelke, M., and Seelow, D. (2014). MutationTaster2: mutation prediction for the deep-sequencing age. Nat Methods 11, 361-362.

62. Cingolani, P., Platts, A., Wang le, L., Coon, M., Nguyen, T., Wang, L., Land, S.J., Lu, X., and Ruden, D.M. (2012). A program for annotating and predicting the effects of single nucleotide polymorphisms, SnpEff: SNPs in the genome of Drosophila melanogaster strain w1118; iso-2; iso-3. Fly (Austin) 6, 80-92.

63. Goecks, J., Nekrutenko, A., Taylor, J., and Galaxy, T. (2010). Galaxy: a comprehensive approach for supporting accessible, reproducible, and transparent computational research in the life sciences. Genome Biol 11, R86.

64. Durbin, R.M., Abecasis, G.R., Altshuler, D.L., Auton, A., Brooks, L.D., Gibbs, R.A., Hurles, M.E., and McVean, G.A. (2010). A map of human genome variation from population-scale sequencing. Nature 467, 1061-1073.

65. Landrum, M.J., Lee, J.M., Riley, G.R., Jang, W., Rubinstein, W.S., Church, D.M., and Maglott, D.R. (2014). ClinVar: public archive of relationships among sequence variation and human phenotype. Nucleic Acids Res 42, D980-985.

66. Ellis, K.L., Frampton, C.M., Pilbrow, A.P., Troughton, R.W., Doughty, R.N., Whalley, G.A., Ellis, C.J., Skelton, L., Thomson, J., Yandle, T.G., et al. (2011). Genomic risk variants at 1p13.3, 1q41, and 3q22.3 are associated with subsequent cardiovascular outcomes in healthy controls and in established coronary artery disease. Circ Cardiovasc Genet 4, 636-646.

67. Purcell, S., Neale, B., Todd-Brown, K., Thomas, L., Ferreira, M.A., Bender, D., Maller, J., Sklar, P., de Bakker, P.I., Daly, M.J., et al. (2007). PLINK: a tool set for whole-genome association and population-based linkage analyses. Am J Hum Genet 81, 559-575.

68. Team, R.C. (2016). R: A language and environment for statistical computing. (R Foundation for Statistical Computing, Vienna, Austria).

69. Benjamini, Y., and Yekutieli, D. (2001). The control of the false discovery rate in multiple testing under dependency. Ann Stat 29, 1165-1188.

70. Neill, N.J., Torchia, B.S., Bejjani, B.A., Shaffer, L.G., and Ballif, B.C. (2010). Comparative analysis of copy number detection by whole-genome BAC and oligonucleotide array CGH. Molecular cytogenetics 3, 11.

71. Firth, H.V., Richards, S.M., Bevan, A.P., Clayton, S., Corpas, M., Rajan, D., Van Vooren, S., Moreau, Y., Pettett, R.M., and Carter, N.P. (2009). DECIPHER: Database of Chromosomal Imbalance and Phenotype in Humans Using Ensembl Resources. Am J Hum Genet 84, 524-533.

72. MacDonald, J.R., Ziman, R., Yuen, R.K., Feuk, L., and Scherer, S.W. (2014). The Database of Genomic Variants: a curated collection of structural variation in the human genome. Nucleic Acids Res 42, D986-992.

73. Kearney, H.M., Thorland, E.C., Brown, K.K., Quintero-Rivera, F., South, S.T., and Working Group of the American College of Medical Genetics Laboratory Quality Assurance, C. (2011). American College of Medical Genetics standards and guidelines for interpretation and reporting of postnatal constitutional copy number variants. Genet Med 13, 680-685.

74. Riggs, E.R., Church, D.M., Hanson, K., Horner, V.L., Kaminsky, E.B., Kuhn, R.M., Wain, K.E., Williams, E.S., Aradhya, S., Kearney, H.M., et al. (2012). Towards an evidence-based process for the clinical interpretation of copy number variation. Clin Genet 81, 403-412.

75. Hanemaaijer, N.M., Sikkema-Raddatz, B., van der Vries, G., Dijkhuizen, T., Hordijk, R., van Essen, A.J., Veenstra-Knol, H.E., Kerstjens-Frederikse, W.S., Herkert, J.C., Gerkes, E.H., et al. (2012). Practical guidelines for interpreting copy number gains detected by high-resolution array in routine diagnostics. Eur J Hum Genet 20, 161-165.

76. Breckpot, J., Thienpont, B., Arens, Y., Tranchevent, L.C., Vermeesch, J.R., Moreau, Y., Gewillig, M., and Devriendt, K. (2011). Challenges of interpreting copy number variation in syndromic and non-syndromic congenital heart defects. Cytogenet Genome Res 135, 251-259.

77. Mills, R.E., Walter, K., Stewart, C., Handsaker, R.E., Chen, K., Alkan, C., Abyzov, A., Yoon, S.C., Ye, K., Cheetham, R.K., et al. (2011). Mapping copy number variation by population-scale genome sequencing. Nature 470, 59-65.

78. Cooper, G.M., Coe, B.P., Girirajan, S., Rosenfeld, J.A., Vu, T.H., Baker, C., Williams, C., Stalker, H., Hamid, R., Hannig, V., et al. (2011). A copy number variation morbidity map of developmental delay. Nat Genet 43, 838-846.

79. Coe, B.P., Witherspoon, K., Rosenfeld, J.A., van Bon, B.W., Vulto-van Silfhout, A.T., Bosco, P., Friend, K.L., Baker, C., Buono, S., Vissers, L.E., et al. (2014). Refining analyses of copy number variation identifies specific genes associated with developmental delay. Nat Genet 46, 1063-1071.

80. Shaikh, T.H., Gai, X., Perin, J.C., Glessner, J.T., Xie, H., Murphy, K., O’Hara, R., Casalunovo, T., Conlin, L.K., D’Arcy, M., et al. (2009). High-resolution mapping and analysis of copy number variations in the human genome: a data resource for clinical and research applications. Genome Res 19, 1682-1690.

81. Vogler, C., Gschwind, L., Rothlisberger, B., Huber, A., Filges, I., Miny, P., Auschra, B., Stetak, A., Demougin, P., Vukojevic, V., et al. (2010). Microarray-based maps of copy-number variant regions in European and sub-Saharan populations. PLoS One 5, e15246.

82. Kircher, M., Witten, D.M., Jain, P., O’Roak, B.J., Cooper, G.M., and Shendure, J. (2014). A general framework for estimating the relative pathogenicity of human genetic variants. Nat Genet 46, 310-315.

83. Picardi, E., and Pesole, G. (2012). Mitochondrial genomes gleaned from human whole-exome sequencing. Nat Methods 9, 523-524.

84. Chua, E.W., Cree, S., Barclay, M.L., Doudney, K., Lehnert, K., Aitchison, A., and Kennedy, M.A. (2015). Exome sequencing and array-based comparative genomic hybridisation analysis of preferential 6-methylmercaptopurine producers. Pharmacogenomics J In press.

85. Li, Y.I., Sanchez-Pulido, L., Haerty, W., and Ponting, C.P. (2015). RBFOX and PTBP1 proteins regulate the alternative splicing of micro-exons in human brain transcripts. Genome Res 25, 1-13.

86. Irimia, M., Weatheritt, R.J., Ellis, J.D., Parikshak, N.N., Gonatopoulos-Pournatzis, T., Babor, M., Quesnel-Vallieres, M., Tapial, J., Raj, B., O’Hanlon, D., et al. (2014). A highly conserved program of neuronal microexons is misregulated in autistic brains. Cell 159, 1511-1523.

87. Zhao, W.W. (2013). Intragenic deletion of RBFOX1 associated with neurodevelopmental/neuropsychiatric disorders and possibly other clinical presentations. Molecular cytogenetics 6, 26.

88. Lale, S., Yu, S., and Ahmed, A. (2011). Complex congenital heart defects in association with maternal diabetes and partial deletion of the A2BP1 gene. Fetal Pediatr Pathol 30, 161-166.

89. Lal, D., Trucks, H., Moller, R.S., Hjalgrim, H., Koeleman, B.P., de Kovel, C.G., Visscher, F., Weber, Y.G., Lerche, H., Becker, F., et al. (2013). Rare exonic deletions of the RBFOX1 gene increase risk of idiopathic generalized epilepsy. Epilepsia 54, 265-271.

90. Gao, C., Ren, S., Lee, J.H., Qiu, J., Chapski, D.J., Rau, C.D., Zhou, Y., Abdellatif, M., Nakano, A., Vondriska, T.M., et al. (2016). RBFox1-mediated RNA splicing regulates cardiac hypertrophy and heart failure. J Clin Invest 126, 195-206.

91. Earle, N.J., Poppe, K.K., Pilbrow, A.P., Cameron, V.A., Troughton, R.W., Skinner, J.R., Love, D.R., Shelling, A.N., Whalley, G.A., Ellis, C.J., et al. (2015). Genetic markers of repolarization and arrhythmic events after acute coronary syndromes. Am Heart J 169, 579-586 e573.

92. Arking, D.E., Reinier, K., Post, W., Jui, J., Hilton, G., O’Connor, A., Prineas, R.J., Boerwinkle, E., Psaty, B.M., Tomaselli, G.F., et al. (2010). Genome-wide association study identifies GPC5 as a novel genetic locus protective against sudden cardiac arrest. PLoS One 5, e9879.

93. Rosenberg, R.D., Shworak, N.W., Liu, J., Schwartz, J.J., and Zhang, L. (1997). Heparan sulfate proteoglycans of the cardiovascular system. Specific structures emerge but how is synthesis regulated? J Clin Invest 100, S67-75.

94. Gunteski-Hamblin, A.M., Song, G., Walsh, R.A., Frenzke, M., Boivin, G.P., Dorn, G. W., 2nd, Kaetzel, M.A., Horseman, N.D., and Dedman, J.R. (1996). Annexin VI overexpression targeted to heart alters cardiomyocyte function in transgenic mice. Am J Physiol 270, H1091-1100.

95. Song, G., Harding, S.E., Duchen, M.R., Tunwell, R., O’Gara, P., Hawkins, T.E., and Moss, S.E. (2002). Altered mechanical properties and intracellular calcium signaling in cardiomyocytes from annexin 6 null-mutant mice. FASEB J 16, 622-624.

96. Camors, E., Monceau, V., and Charlemagne, D. (2005). Annexins and Ca2+ handling in the heart. Cardiovasc Res 65, 793-802.

97. Meadows, S.M., and Cleaver, O. (2015). Annexin A3 Regulates Early Blood Vessel Formation. PLoS One 10, e0132580.

98. Adhikari, N., Guan, W., Capaldo, B., Mackey, A.J., Carlson, M., Ramakrishnan, S., Walek, D., Gupta, M., Mitchell, A., Eckman, P., et al. (2014). Identification of a new target of miR-16, Vacuolar Protein Sorting 4a. PLoS One 9, e101509.

99. Porrello, E.R., Johnson, B.A., Aurora, A.B., Simpson, E., Nam, Y.J., Matkovich, S.J., Dorn, G.W., 2nd, van Rooij, E., and Olson, E.N. (2011). MiR-15 family regulates postnatal mitotic arrest of cardiomyocytes. Circ Res 109, 670-679.

100. Hullinger, T.G., Montgomery, R.L., Seto, A.G., Dickinson, B.A., Semus, H.M., Lynch, J.M., Dalby, C.M., Robinson, K., Stack, C., Latimer, P.A., et al. (2012). Inhibition of miR-15 protects against cardiac ischemic injury. Circ Res 110, 71-81.

101. Venkateswaran, A., Sekhar, K.R., Levic, D.S., Melville, D.B., Clark, T.A., Rybski, W.M., Walsh, A.J., Skala, M.C., Crooks, P.A., Knapik, E.W., et al. (2014). The NADH oxidase ENOX1, a critical mediator of endothelial cell radiosensitization, is crucial for vascular development. Cancer Res 74, 38-43.

102. Weng, L., Hubner, R., Claessens, A., Smits, P., Wauters, J., Tylzanowski, P., Van Marck, E., and Merregaert, J. (2003). Isolation and characterization of chondrolectin (Chodl), a novel C-type lectin predominantly expressed in muscle cells. Gene 308, 21-29.

103. Coppinger, J., McDonald-McGinn, D., Zackai, E., Shane, K., Atkin, J.F., Asamoah, A., Leland, R., Weaver, D.D., Lansky-Shafer, S., Schmidt, K., et al. (2009). Identification of familial and de novo microduplications of 22q11.21-q11.23 distal to the 22q11.21 microdeletion syndrome region. Hum Mol Genet 18, 1377-1383.

104. Wentzel, C., Fernstrom, M., Ohrner, Y., Anneren, G., and Thuresson, A.C. (2008). Clinical variability of the 22q11.2 duplication syndrome. Eur J Med Genet 51, 501-510.

105. Coe, B.P., Witherspoon, K., Rosenfeld, J.A., van Bon, B.W., Vulto-van Silfhout, A.T., Bosco, P., Friend, K.L., Baker, C., Buono, S., Vissers, L.E., et al. (2014). Refining analyses of copy number variation identifies specific genes associated with developmental delay. Nat Genet 46, 1063-1071.

106. Pipes, L., Li, S., Bozinoski, M., Palermo, R., Peng, X., Blood, P., Kelly, S., Weiss, J.M., Thierry-Mieg, J., Thierry-Mieg, D., et al. (2013). The non-human primate reference transcriptome resource (NHPRTR) for comparative functional genomics. Nucleic Acids Res 41, D906-914.

107. Vadas, O., Dbouk, H.A., Shymanets, A., Perisic, O., Burke, J.E., Abi Saab, W.F., Khalil, B.D., Harteneck, C., Bresnick, A.R., Nurnberg, B., et al. (2013). Molecular determinants of PI3Kgamma-mediated activation downstream of G-protein-coupled receptors (GPCRs). Proc Natl Acad Sci U S A 110, 18862-18867.

108. Izumi, Y. (2013). Drug-induced takotsubo cardiomyopathy. Heart Fail Clin 9, 225-231, ix-x.

109. Ueyama, T. (2004). Emotional stress-induced Tako-tsubo cardiomyopathy: animal model and molecular mechanism. Ann N Y Acad Sci 1018, 437-444.

110. Kido, K., and Guglin, M. (2017). Drug-Induced Takotsubo Cardiomyopathy. Journal of cardiovascular pharmacology and therapeutics, 1074248417708618.

111. Schreyer, S., Ledwig, D., Rakatzi, I., Kloting, I., and Eckel, J. (2003). Insulin receptor substrate-4 is expressed in muscle tissue without acting as a substrate for the insulin receptor. Endocrinology 144, 1211-1218.

112. Fantin, V.R., Lavan, B.E., Wang, Q., Jenkins, N.A., Gilbert, D.J., Copeland, N.G., Keller, S.R., and Lienhard, G.E. (1999). Cloning, tissue expression, and chromosomal location of the mouse insulin receptor substrate 4 gene. Endocrinology 140, 1329-1337.

113. Mertens, F., Moller, E., Mandahl, N., Picci, P., Perez-Atayde, A.R., Samson, I., Sciot, R., and Debiec-Rychter, M. (2011). The t(X;6) in subungual exostosis results in transcriptional deregulation of the gene for insulin receptor substrate 4. Int J Cancer 128, 487-491.

114. Duenez-Guzman, E.A., and Haig, D. (2014). The evolution of reproduction-related NLRP genes. J Mol Evol 78, 194-201.

115. Snoep, J.D., Gaussem, P., Eikenboom, J.C., Emmerich, J., Zwaginga, J.J., Holmes, C. E., Vos, H.L., de Groot, P.G., Herrington, D.M., Bray, P.F., et al. (2010). The minor allele of GP6 T13254C is associated with decreased platelet activation and a reduced risk of recurrent cardiovascular events and mortality: results from the SMILE-Platelets project. J Thromb Haemost 8, 2377-2384.

116. Bray, P.F., Howard, T.D., Vittinghoff, E., Sane, D.C., and Herrington, D.M. (2007). Effect of genetic variations in platelet glycoproteins Ibalpha and VI on the risk for coronary heart disease events in postmenopausal women taking hormone therapy. Blood 109, 1862-1869.

117. Lorenz, K., Schmitt, J.P., Schmitteckert, E.M., and Lohse, M.J. (2009). A new type of ERK1/2 autophosphorylation causes cardiac hypertrophy. Nat Med 15, 75-83.

118. Dimassi, S., Labalme, A., Lesca, G., Rudolf, G., Bruneau, N., Hirsch, E., Arzimanoglou, A., Motte, J., de Saint Martin, A., Boutry-Kryza, N., et al. (2014). A subset of genomic alterations detected in rolandic epilepsies contains candidate or known epilepsy genes including GRIN2A and PRRT2. Epilepsia 55, 370-378.

119. Zhao, Y.Y., Sawyer, D.R., Baliga, R.R., Opel, D.J., Han, X., Marchionni, M.A., and Kelly, R.A. (1998). Neuregulins promote survival and growth of cardiac myocytes. Persistence of ErbB2 and ErbB4 expression in neonatal and adult ventricular myocytes. J Biol Chem 273, 10261-10269.

120. Berg, R.W., Leung, E., Gough, S., Morris, C., Yao, W.P., Wang, S.X., Ni, J., and Krissansen, G.W. (1999). Cloning and characterization of a novel beta integrin-related cDNA coding for the protein TIED (“ten beta integrin EGF-like repeat domains”) that maps to chromosome band 13q33: A divergent stand-alone integrin stalk structure. Genomics 56, 169-178.

121. Wheeler, H.E., Shah, K.P., Brenner, J., Garcia, T., Aquino-Michaels, K., Consortium, G.T., Cox, N.J., Nicolae, D.L., and Im, H.K. (2016). Survey of the Heritability and Sparse Architecture of Gene Expression Traits across Human Tissues. PLoS Genet 12, e1006423.

122. Wittstein, I.S., Thiemann, D.R., Lima, J.A., Baughman, K.L., Schulman, S.P., Gerstenblith, G., Wu, K.C., Rade, J.J., Bivalacqua, T.J., and Champion, H.C. (2005). Neurohumoral features of myocardial stunning due to sudden emotional stress. N Engl J Med 352, 539-548.

123. Voight, B.F., Kang, H.M., Ding, J., Palmer, C.D., Sidore, C., Chines, P.S., Burtt, N.P., Fuchsberger, C., Li, Y., Erdmann, J., et al. (2012). The metabochip, a custom genotyping array for genetic studies of metabolic, cardiovascular, and anthropometric traits. PLoS Genet 8, e1002793.

124. Eitel, I., Moeller, C., Munz, M., Stiermaier, T., Meitinger, T., Thiele, H., and Erdmann, J. (2017). Genome-wide association study in takotsubo syndrome - Preliminary results and future directions. Int J Cardiol 236, 335-339.

125. Redon, R., Ishikawa, S., Fitch, K.R., Feuk, L., Perry, G.H., Andrews, T.D., Fiegler, H., Shapero, M.H., Carson, A.R., Chen, W., et al. (2006). Global variation in copy number in the human genome. Nature 444, 444-454.

126. McCarroll, S.A., and Altshuler, D.M. (2007). Copy-number variation and association studies of human disease. Nat Genet 39, S37-42.

127. Stranger, B.E., Forrest, M.S., Dunning, M., Ingle, C.E., Beazley, C., Thorne, N., Redon, R., Bird, C.P., de Grassi, A., Lee, C., et al. (2007). Relative impact of nucleotide and copy number variation on gene expression phenotypes. Science 315, 848-853.

128. Purcell, S.M., Moran, J.L., Fromer, M., Ruderfer, D., Solovieff, N., Roussos, P., O’Dushlaine, C., Chambert, K., Bergen, S.E., Kahler, A., et al. (2014). A polygenic burden of rare disruptive mutations in schizophrenia. Nature 506, 185-190.

129. Helbig, I., Swinkels, M.E., Aten, E., Caliebe, A., van’t Slot, R., Boor, R., von Spiczak, S., Muhle, H., Jahn, J.A., van Binsbergen, E., et al. (2014). Structural genomic variation in childhood epilepsies with complex phenotypes. Eur J Hum Genet 22, 896-901.

130. Hooli, B.V., Kovacs-Vajna, Z.M., Mullin, K., Blumenthal, M.A., Mattheisen, M., Zhang, C., Lange, C., Mohapatra, G., Bertram, L., and Tanzi, R.E. (2014). Rare autosomal copy number variations in early-onset familial Alzheimer’s disease. Mol Psychiatry 19, 676-681.

131. Gilissen, C., Hehir-Kwa, J.Y., Thung, D.T., van de Vorst, M., van Bon, B.W., Willemsen, M.H., Kwint, M., Janssen, I.M., Hoischen, A., Schenck, A., et al. (2014). Genome sequencing identifies major causes of severe intellectual disability. Nature 511, 344-347.

132. Mefford, H.C., Yendle, S.C., Hsu, C., Cook, J., Geraghty, E., McMahon, J.M., Eeg-Olofsson, O., Sadleir, L.G., Gill, D., Ben-Zeev, B., et al. (2011). Rare copy number variants are an important cause of epileptic encephalopathies. Ann Neurol 70, 974-985.

133. Hitz, M.P., Lemieux-Perreault, L.P., Marshall, C., Feroz-Zada, Y., Davies, R., Yang, S.W., Lionel, A.C., D’Amours, G., Lemyre, E., Cullum, R., et al. (2012). Rare copy number variants contribute to congenital left-sided heart disease. PLoS Genet 8, e1002903.

134. White, P.S., Xie, H.M., Werner, P., Glessner, J., Latney, B., Hakonarson, H., and Goldmuntz, E. (2014). Analysis of chromosomal structural variation in patients with congenital left-sided cardiac lesions. Birth Defects Res A Clin Mol Teratol.

135. Fakhro, K.A., Choi, M., Ware, S.M., Belmont, J.W., Towbin, J.A., Lifton, R.P., Khokha, M.K., and Brueckner, M. (2011). Rare copy number variations in congenital heart disease patients identify unique genes in left-right patterning. Proc Natl Acad Sci U S A 108, 2915-2920.

136. Glessner, J.T., Bick, A.G., Ito, K., Homsy, J.G., Rodriguez-Murillo, L., Fromer, M., Mazaika, E., Vardarajan, B., Italia, M., Leipzig, J., et al. (2014). Increased frequency of de novo copy number variants in congenital heart disease by integrative analysis of single nucleotide polymorphism array and exome sequence data. Circ Res 115, 884-896.

137. Barc, J., Briec, F., Schmitt, S., Kyndt, F., Le Cunff, M., Baron, E., Vieyres, C., Sacher, F., Redon, R., Le Caignec, C., et al. (2011). Screening for copy number variation in genes associated with the long QT syndrome: clinical relevance. J Am Coll Cardiol 57, 40-47.

138. Silversides, C.K., Lionel, A.C., Costain, G., Merico, D., Migita, O., Liu, B., Yuen, T., Rickaby, J., Thiruvahindrapuram, B., Marshall, C.R., et al. (2012). Rare copy number variations in adults with tetralogy of Fallot implicate novel risk gene pathways. PLoS Genet 8, e1002843.

139. Roberts, J.L., Hovanes, K., Dasouki, M., Manzardo, A.M., and Butler, M.G. (2014). Chromosomal microarray analysis of consecutive individuals with autism spectrum disorders or learning disability presenting for genetic services. Gene 535, 70-78.

140. Battaglia, A., Doccini, V., Bernardini, L., Novelli, A., Loddo, S., Capalbo, A., Filippi, T., and Carey, J.C. (2013). Confirmation of chromosomal microarray as a first-tier clinical diagnostic test for individuals with developmental delay, intellectual disability, autism spectrum disorders and dysmorphic features. Eur J Paediatr Neurol 17, 589-599.

141. Chong, W.W., Lo, I.F., Lam, S.T., Wang, C.C., Luk, H.M., Leung, T.Y., and Choy, K.W. (2014). Performance of chromosomal microarray for patients with intellectual disabilities/developmental delay, autism, and multiple congenital anomalies in a Chinese cohort. Molecular cytogenetics 7, 34.

142. Mc Cormack, A., Claxton, K., Ashton, F., Asquith, P., Atack, E., Mazzaschi, R., Moverley, P., O’Connor, R., Qorri, M., Sheath, K., et al. (2016). Microarray testing in clinical diagnosis: an analysis of 5,300 New Zealand patients. Molecular cytogenetics 9, 29.

143. Ruusala, A., and Aspenstrom, P. (2004). Isolation and characterisation of DOCK8, a member of the DOCK180-related regulators of cell morphology. FEBS Lett 572, 159-166.

144. Engelhardt, K.R., McGhee, S., Winkler, S., Sassi, A., Woellner, C., Lopez-Herrera, G., Chen, A., Kim, H.S., Lloret, M.G., Schulze, I., et al. (2009). Large deletions and point mutations involving the dedicator of cytokinesis 8 (DOCK8) in the autosomal-recessive form of hyper-IgE syndrome. J Allergy Clin Immunol 124, 1289-1302 e1284.

145. Al Mutairi, M., Al-Mousa, H., AlSaud, B., Hawwari, A., AlJoufan, M., AlWesaibi, A., AlHalees, Z., and Al-Mayouf, S.M. (2014). Grave aortic aneurysmal dilatation in DOCK8 deficiency. Mod Rheumatol 24, 690-693.

146. Sterneckert, J.L., Reinhardt, P., and Scholer, H.R. (2014). Investigating human disease using stem cell models. Nat Rev Genet 15, 625-639.

147. Ma, D., Wei, H., Lu, J., Ho, S., Zhang, G., Sun, X., Oh, Y., Tan, S.H., Ng, M.L., Shim, W., et al. (2013). Generation of patient-specific induced pluripotent stem cell-derived cardiomyocytes as a cellular model of arrhythmogenic right ventricular cardiomyopathy. Eur Heart J 34, 1122-1133.

